# Multidrug resistant *E. coli* isolate along with ESBL production and Antimicrobial Susceptibility pattern from patients of Urinary Tract Infection from tertiary care hospital, Delhi

**DOI:** 10.1101/2025.09.05.674450

**Authors:** Thakur Datt, Shyama Datt, Simrita Singh, N. P. Singh

## Abstract

**Background:** Third generation Cephalosporins resistances among *Escherichia coli* due to production of ESBL, mainly the bla-CTX-M, bla-TEM and bla-SHV genes, poses serious challenges in the clinical utility of these drugs in healthcare settings. This study was undertaken to detect prevalence of ESBL producing MDR *E. coli* isolates by various phenotypic and molecular method from in or out patients with urinary tract infection (UTI).

**Methods:** A total of 56 non-repetitive isolates of *E. coli* from in or out patients of UTI were subjected to antimicrobial susceptibility testing. Screening for the production of extended-spectrum β-lactamases (ESBL) was determined in third generation Cephalosporins resistant isolates by minimum inhibitory concentration (MIC) E-strip, followed by phenotypic confirmation by DDST, CDST and ESBL E-strip method. Further confirmation was done by molecular method, PCR was carried out for detection of bla-CTX-M, bla-TEM and bla-SHV.

**Results:** All strains (56/56) were observed as ESBL producers by MIC E-strip method, 94.64% (53/56) isolates were observed as ESBL producer by DDST and 67.86% (38/56) were observed by CDST. The overall ESBL production among all MDR *E. coli* isolates was 89.29% (50/56) which includes any one of the three genes. The prevalence of bla-CTX-M (80.36%, 45/56) was highest, followed by bla-TEM (50%, 28/56) and bla-SHV (10.71%, 6/56).

**Conclusion:** In our study we found bla-CTX-M gene is predominantly circulating among all three ESBL genes in the clinical isolates of MDR *E. coli* that were producers of ESBL.

## Introduction

Multidrug resistance (MDR) in among pathogenic *Escherichia coli* (*E. coli*) is one of the leading cause of increased mortality and morbidity, in both developed and developing countries. *E. coli* is a common commensal, so it can easily colonize urinary tract and serves as a most common cause of Urinary tract infection (UTI) and bacteremia in humans. (1-Paterson 2005), with an estimated 150 million UTIs per annum worldwide (2-Rahman 2009, 3-Francesco 2007, 4-Martins 2010). Although UTI is treatable, but due to antimicrobial resistance (AMR) in the Enterobacteriaceae family, particularly in *E. coli* is now increasingly becoming tough to control (1-Paterson 2005). Now these organisms posing significant social and economic burden on the community and public health departments (5-Mizrahi 2021).

The early treatment of UTI includes identification of causative agent, antimicrobial resistance pattern and administration of appropriate empirical therapy, which simultaneously decreases the rate of morbidity [3-Francesco 2007, 6-Wagenlehner 2004]. However, antimicrobial therapy has significantly contributed to the management of the infection, but misuse of antimicrobials has led to an increase of the MDR and consequently spread of resistance by horizontal gene transfer (7-Arjunan 2010, 2-Rahman 2009, 8-Igwe 2013).

β-lactamase production has been known as the main mechanism for β-lactam drugs inactivation, due to the presence of several types of enzymes, including ESBLs among *Enterobactericeae* (1-Paterson 2005). The ESBLs that hydrolyse oxyimino β-lactams like Ceftazidime, Cefotaxime, Ceftriaxone and Monobactum but have no effect on the Cephamycins and Carbapenem, have emerged as an important mechanism drug resistance amongst uropathogens (9-Ramazanzadeh 2010). Additionally, these strains are often resistant to other classes of antimicrobial agents, such as Fluoroquinolones, Trimethoprime, Sulfamethoxazole and Aminoglycosides (10-Park 2012, Qureshi 2012).

Therefore, it is wise to know about antibiotic resistance pattern, so that the selection of the antimicrobial agent for the treatment should be obstinate, on the basis of its real time susceptibility pattern. Thus, the knowledge of local antimicrobial susceptibility pattern for *E. coli* is essential for providing an accurate empirical therapy. So, the main objective of this study was to evaluate the AMR pattern and possible ESBL producer among MDR *E. coli* strains isolated form urine samples of patients of UTI form Delhi.

## Materials & Methods

### Samples

This study was carried out in the Department of Microbiology on total of 56 non-repetitive urinary MDR *E. coli* isolates, isolated from clinically suspected cases of UTI admitted in hospital or outpatient department of Guru Teg Bahadur Hospital, a tertiary care hospital in Delhi, between November 2017 to October 2020.

### Sample collection

Freshly voided, midstream urine was collected directly into the sterile wide mouth container. For each patient the collection date, age, sex, urine culture result, identification of the bacterial strain and the corresponding AST results were registered.

### Study population

Patients of age ≥18 to 80 years with atleast one sign or symptom of UTI (dysuria, frequency, urgency, perineal pain, flank pain or costovertebral tenderness) were included in this study.

### Exclusion criteria

Patient’s age <18 years, inpatient admission a week prior to presentation in OPD, antibiotics usage within a week and polymorphic bacterial growth (11-Wilson ML 2004).

Written informed consent was obtained from all patients. The study was conducted after due approval from Institutional Ethics Committee-Human Research of UCMS & GTBH, Delhi.

### Urine culture

Urine sample was processed and streaked on Cystein Lactose Electrolyte Deficient (CLED) agar medium using a calibrated loop of 4 mm diameter. The culture plates were incubated at 37°C for overnight and monomorphic yellow colonies with 3-4 mm diameter were looked out. Count was expressed as colony forming units (cfu) per millimeter (ml). The urine cultures were classified as negative when bacteria growth was lower than 10^3^ cfu/ml and positive when monomorphic bacterial growth with significant bacteriuria (≥10^5^ cfu/ml) (12-Kass 1957).

### Identification

Bacterial isolates were confirmed as *E. coli* on the basis of colonial morphology, microscopy and by using a battery of biochemical tests (13-Collee JG, 15-Miles RS, Watt B 2006).

### Antimicrobial susceptibility test

The AST was performed using the modified Kirby-Bauer disk diffusion method (14-CLSI 2020, 15-Bauer et al 1966). Commercially procured antimicrobial-impregnated disks (Himedia, Mumbai, India) were placed onto the surface of Mueller-Hinton Agar, the 24 antibiotics viz. Imipenem (10 mcg), Meropenem (10 mcg), Ertapenem (10 mcg), Cefotaxime (30 mcg), Ceftazidime (30 mcg), Ceftraixone (30 mcg), Cefoxitin (30 mcg), Cotrimoxazole (25 mcg), Choramphenicol (30 mcg), Amoxicillin-clavulanic acid (20/10 mcg), Amikacin (30 mcg), Ampicillin (10 mcg), Fosfomycin (200 µg), Gentamicin (10 mcg), Levofloxacin (5 mcg), Nitrofurantoin (300 mcg), Norfloxacin (10 mcg), Nalidixic acid (30 mcg), Netilimicin (30 mcg), Piperacillin+Tazobactm (100/10 mcg), Tetracyclin (30 mcg), Tigecyclin (30 mcg), Tobramycin (10 mcg) and Aztreonam (30 mcg) were tested. The plates were incubated at 37°C overnight in ambient air and result was analyzed according to CLSI guidelines (14-CLSI M100 S30). An isolate was considered as multidrug resistant if it was resistant to ≥3 groups of antibiotics (16-Magiorakos AP, Srinivasan A 2011) and in present study an isolate was considered as MDR if found resistant to any of the third generation Cephalosporines, any of Carbapenems and any other class of antibiotic.

### Minimum Inhibitory Concentration (MIC)

MIC for third generation Cephalosporins was determined by commercially available MIC-strip containing gradient of antimicrobial concentrations of Cefotaxime (CTX) (0.016-256 µg/ml) and Ceftazidime (CAZ) (0.016-256 µg/ml) (Himedia, Mumbai, India). MIC of Cefotaxime and Ceftazidime was determined on the basis of CLSI breakpoint (14-CLSI, 2020).

### Phenotypic screening for ESBL production

The screening for ESBL production was carried on the isolates resistant to either of any third generation Cephalosporin (CAZ, CTX and CTR).

### Combined-Disc Synergy Test (CDST)

Third generation Cephalosporin resistant isolates were subjected to CAZ and Ceftazidime-clavulanic Acid (CAC) combination discs. The presence of ESBL activity was indicated by a ≥5 mm increase in zone diameter for Ceftazidime in combination with clavulanic acid (inhibitor) compared Ceftazidime alone (14-CLSI 2020).

### Double-Disc Synergy Test (DDST)

It was carried out by placing four discs viz Ceftazidime (30 mcg), Cefotaxime (30 mcg), Ceftriaxone (30 mcg), Cefpodoxime (30 mcg) radially at a distance of 20 mm each from a disc containing Amoxicillin-Clavulanic acid (20/10 mcg) on a lawn culture of the MDR *E. coli* isolate on MHA plate (14-CLSI 2020). The plates were incubated at 37°C. The strain was confirmed as ESBL producer, if the zone size around any of the discs were enhanced toward amoxicillin-clavulanic acid disc. *K. pneumonia* ATCC 700603 (ESBL producer) strain was used as positive control (14-CLSI 2020).

### ESBL E-strip

This was carried out for presumptive screening of ESBL using commercially available MIC ESBL test strip (Himedia, Mumbai, India). The range of antimicrobials in ESBL E-strip had Ceftazidime (CAZ), gradient at one end (0.5-32 µg/ml) and gradient of Ceftazidime (1-64 µg/ml) plus clavulanic acid (CAZ^+^) (0.0064-4 µg/ml) at other end. The isolate was considered as ESBL producer if MIC ratio of (CAZ/CAZ^+^) was >8 (14-CLSI 2020).

### Molecular test

#### Genomic DNA isolation

The DNA was isolated as per the Qiagen (Germany), Genomic DNA kit following the manufacturer protocol. The DNA was used to perform PCR for identification of genes viz. bla-CTX-M, bla-SHV and bla-TEM associated with β-lactamase antibiotic resistance (17-Bhattacharjee, Sen, Anupurba 2007). Target gene with primer sequences and molecular weight of amplified DNA generated is given in Table-1.

**Table 1.**
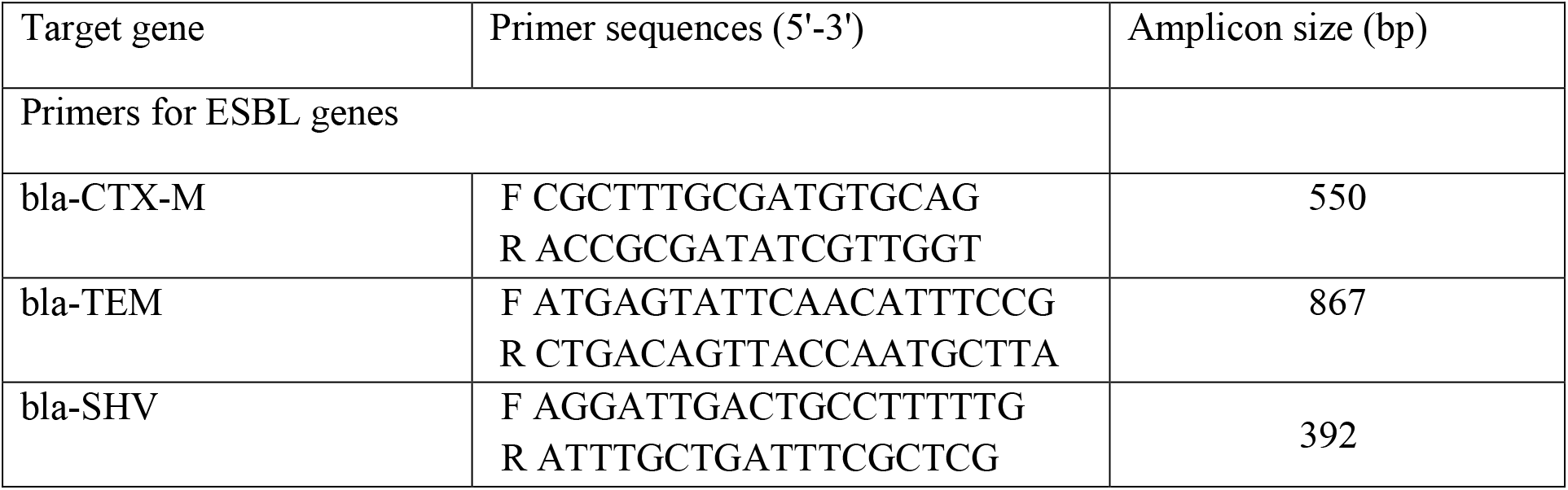
Showing target gene with primer sequences.

#### PCR conditions

Each PCR tube contain, a total volume of 25 µl including 12.5 µl PCR master mix, 1 µl each of forward and reverse primer (10 pmol working concentration), 2 µl of extracted DNA and nuclease free water to make up the volume. The amplification was performed in a Thermal Cycler (ABI system). After an initial denaturation at 95°C for 5 min. the reaction mixed was subjected to 35 cycles of amplification of 40 sec at 95°C, 40 sec at 55°C and 1 min at 72°C, and final extension of 7 min at 72°C.

#### Gel-electrophoresis

PCR product of amplified genes was then separated by gel-electrophoresis on 1.5% agarose gel and stained with ethidium bromide, a molecular weight bands (100 bp DNA ladder, Himedia,

Mumbai, India) was used to determine the size of the amplicons. Documentation was done in gel doc system.

## Results

A total 56 MDR *E. coli* strains isolated from clinically suspected UTI patients were included for this study. The age distribution of the patients in the sample set was 18-80 years (mean & standard deviation 43.18 ± 18.09 years). From total 56 patients, 43/56 (76.79%) were female, and 13/56 (23.20%) male patients, had bacteriuria.

The prevalence of UTI was highest within 21 to 30 years of age group (25%), followed by 31 to 40 years (21.43 %).

Among the female patients most affected age groups were 21 to 30 and 31 to 40 (25.58%) and in the case of male patients most affected age group was 21 to 30 (30.77%) followed by 71 to 80 (23.08%). Age wise and sex wise distribution of UTI patients is shown in figure-1.

**Figure 1.**
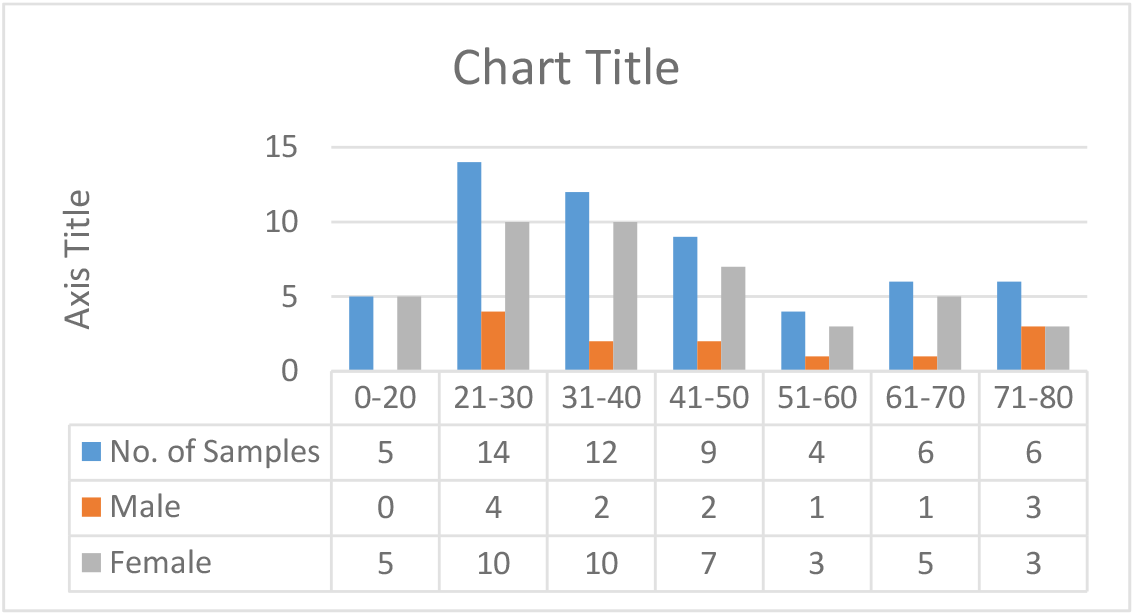
Age wise and sex wise distribution of UTI patients.

Antimicrobial resistance profile among MDR *E. coli* isolates are summarized in Table-2. No resistance has been observed for Fosfomycin, and then followed by Tigecycline (19.64%), Nitrofurantoin (28.57%) and Chloramphenicol (30.36%) respectively. The commonly recommended antimicrobials i.e. Cotrimoxazole and Ampicillin showed high resistant rates i.e. 87.5% and 100% respectively. We have observed absolute resistance to Cefotaxime, Ceftriaxone, Ampicillin, Nalidixic acid, Levofloxacin, and Norfloxacin.

**Table 2.**
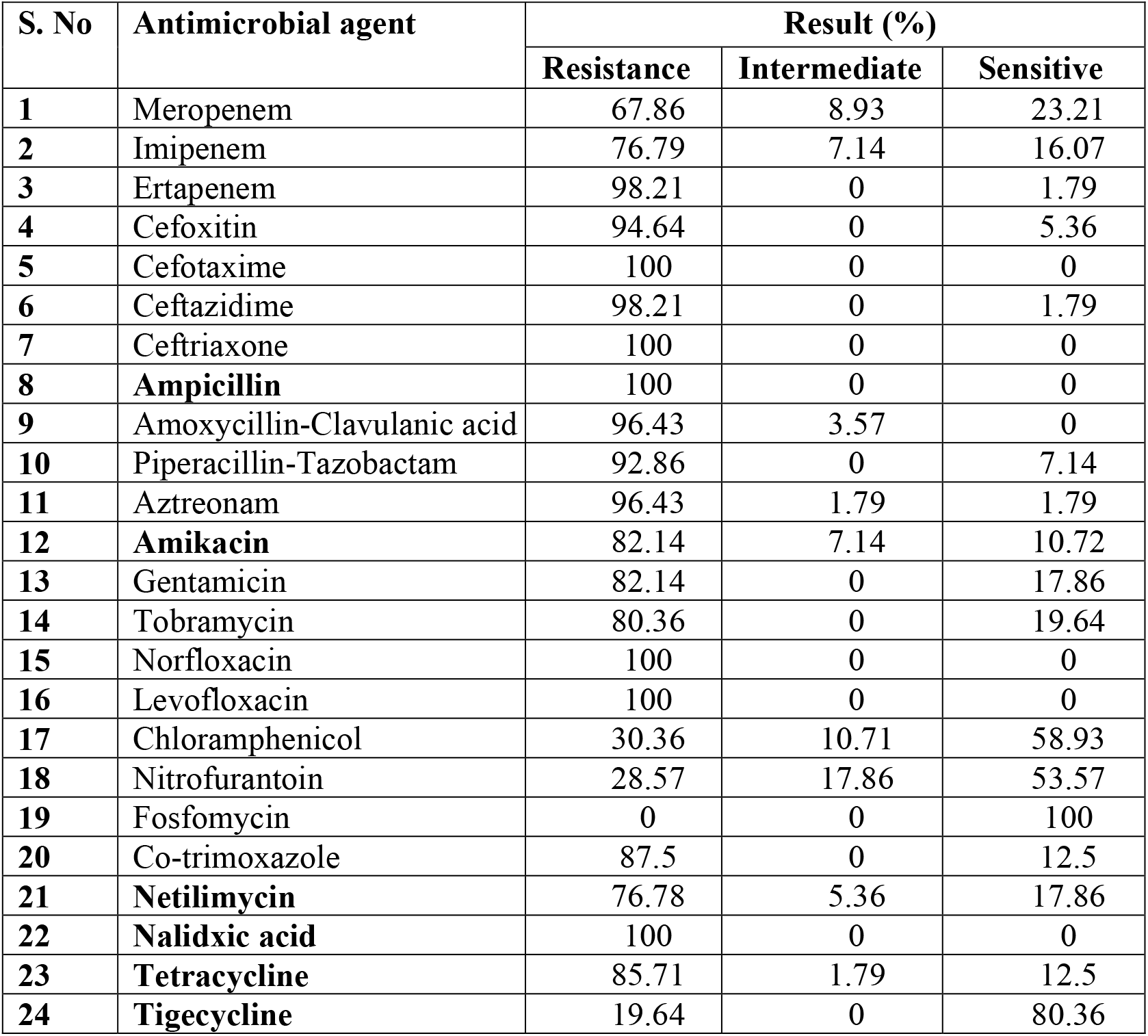
Antimicrobials resistance among UTI patients.

| S. No | Antimicrobial agent | Result (%) |  |  |
| --- | --- | --- | --- | --- |
|  |  | Resistance | Intermediate | Sensitive |
| 1 | Meropenem | 67.86 | 8.93 | 23.21 |
| 2 | Imipenem | 76.79 | 7.14 | 16.07 |
| 3 | Ertapenem | 98.21 | 0 | 1.79 |
| 4 | Cefoxitin | 94.64 | 0 | 5.36 |
| 5 | Cefotaxime | 100 | 0 | 0 |
| 6 | Ceftazidime | 98.21 | 0 | 1.79 |
| 7 | Ceftriaxone | 100 | 0 | 0 |
| 8 | **Ampicillin** | 100 | 0 | 0 |
| 9 | Amoxycillin-Clavulanic acid | 96.43 | 3.57 | 0 |
| 10 | Piperacillin-Tazobactam | 92.86 | 0 | 7.14 |
| 11 | Aztreonam | 96.43 | 1.79 | 1.79 |
| 12 | **Amikacin** | 82.14 | 7.14 | 10.72 |
| 13 | Gentamicin | 82.14 | 0 | 17.86 |
| 14 | Tobramycin | 80.36 | 0 | 19.64 |
| 15 | Norfloxacin | 100 | 0 | 0 |
| 16 | Levofloxacin | 100 | 0 | 0 |
| 17 | Chloramphenicol | 30.36 | 10.71 | 58.93 |
| 18 | Nitrofurantoin | 28.57 | 17.86 | 53.57 |
| 19 | Fosfomycin | 0 | 0 | 100 |
| 20 | Co-trimoxazole | 87.5 | 0 | 12.5 |
| 21 | **Netilmicin** | 76.78 | 5.36 | 17.86 |
| 22 | **Nalidixic acid** | 100 | 0 | 0 |
| 23 | **Tetracycline** | 85.71 | 1.79 | 12.5 |
| 24 | **Tigecycline** | 19.64 | 0 | 80.36 |

Among β-lacatam/β-lacatamase inhibitor drugs, highest resistance was observed with Amoxicillin-Clavulanic acid (96.43%) as compared to Piperacillin-tazobactam (92.86%) (Table-2). Among Aminoglycosides highest resistance was observed with Amikacin (82.14%), Gentamycin (82.14%) followed by Tobramycin (80.36%) and Netillin (76.78%).

Among Quinolones higher resistance was observed in Norfloxacin, Nalidixic acid and Levofloxacin i.e. 100%. Among Carbapenems highest resistance was observed in Ertapenem (98.21%) followed by Imipenem (76.79%) and Meropenem (67.86%). For Cephems including Cephalosporins II and III generations we have observed highest resistance to Cefotaxime (100%), Ceftriaxone (100%), followed by Ceftazidime (98.21%) and Cefoxitin (94.64%).

The resistance pattern for 24 antimicrobial agents of different classes were analyzed and shown in Table-3. 46 different resistance patterns amongst 56 MDR *E. coli* isolates have been observed. One of the patterns of three isolates are pan resistant showing sensitivity to only Fosfomycin and Tigecycline, followed by five isolates showing resistance to 21 antimicrobials showing sensitivity to only Fosfomycin, Tigecycline & Nitrofurantoin. One of the isolate showed resistance to 22 antimicrobials and simultaneously intermediate to one antimicrobial and two isolates resistant to 20 antimicrobials. Furthermore, in 17 isolates we have observed seven different types of patterns and 39 different isolates in 39 different patterns.

**Table 3.**
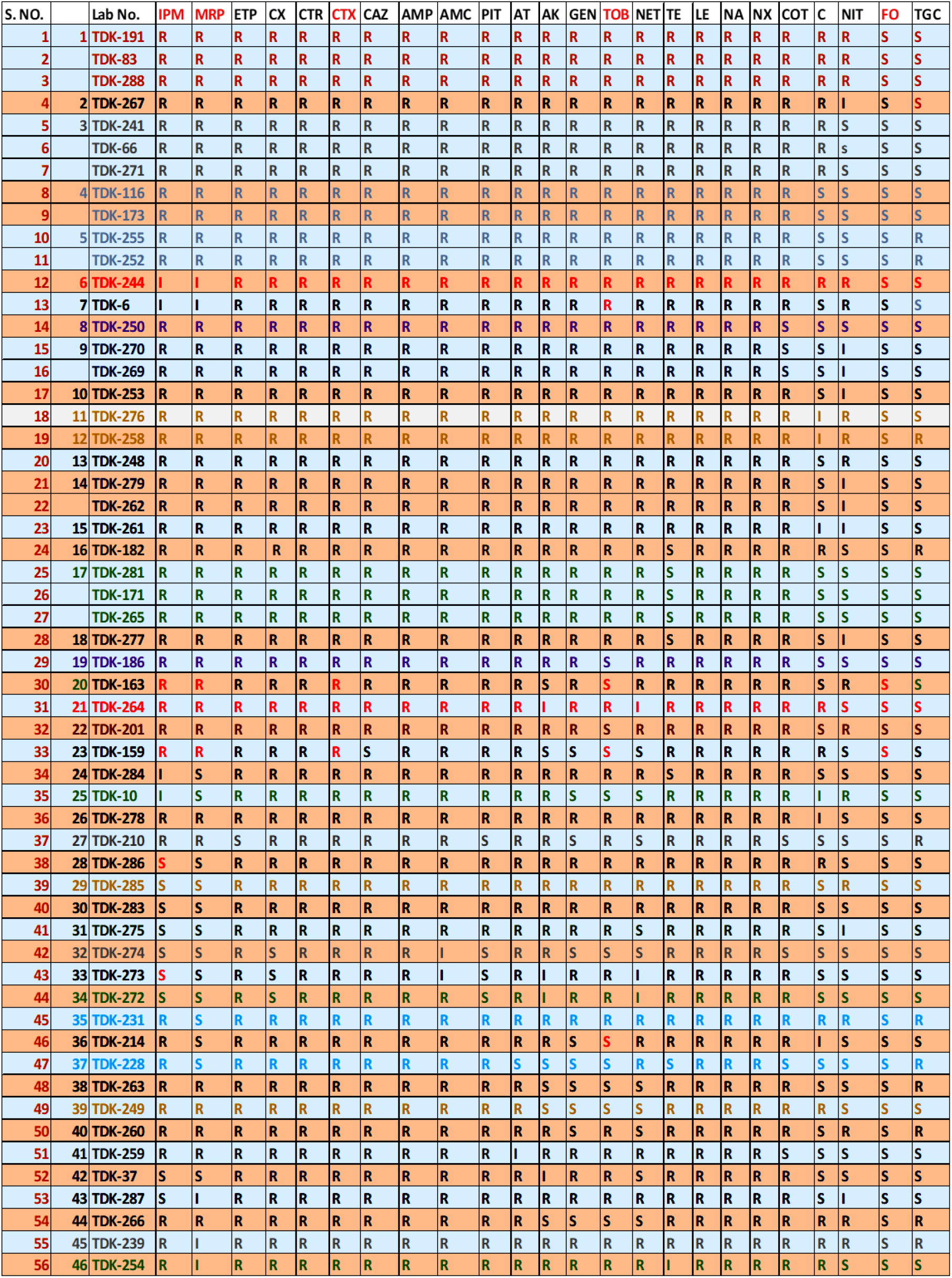
– Showing resistance pattern of 56 MDR *E. coli* isolates isolated from Urine.

ESBL screening was performed on isolates resistant to either of 3^rd^ generation Cephalosporins. MIC values for Cefotaxime and Ceftazidime was evaluated using MIC E-strips. For Cefotaxime all strains were resistant, while one strain was intermediate for Ceftazidime and rest all were resistant.

The mean average value and standard deviated for Cefotaxime and Ceftazidime were 252±29.93 & 247±44.58. All strains (56/56) were observed as ESBL producers by MIC E-strip method, 94.64% (53/56) isolates were observed as ESBL producer by DDST and 67.86% (38/56) were observed by CDST, Figure-2.

**Figure 2.**
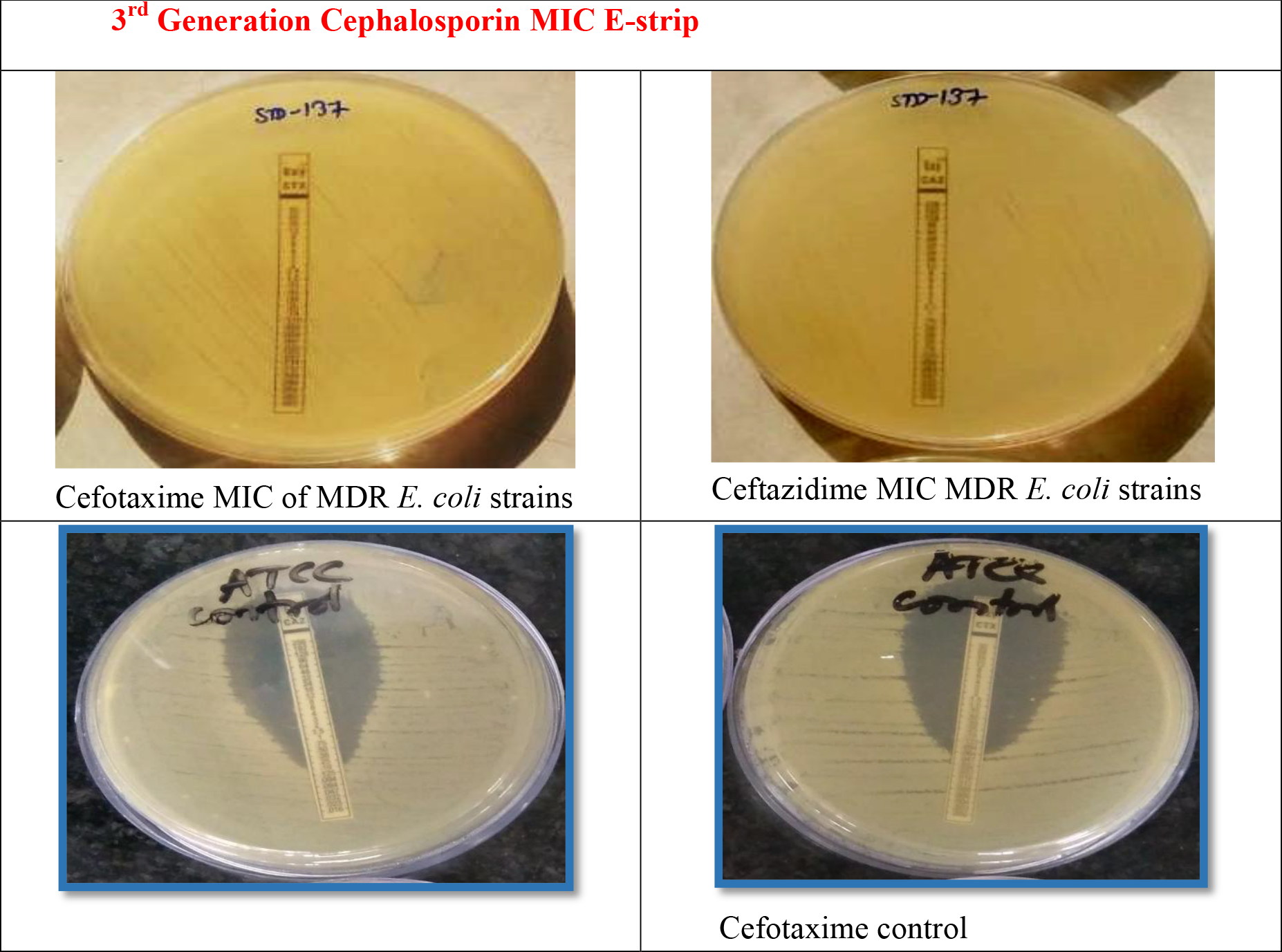
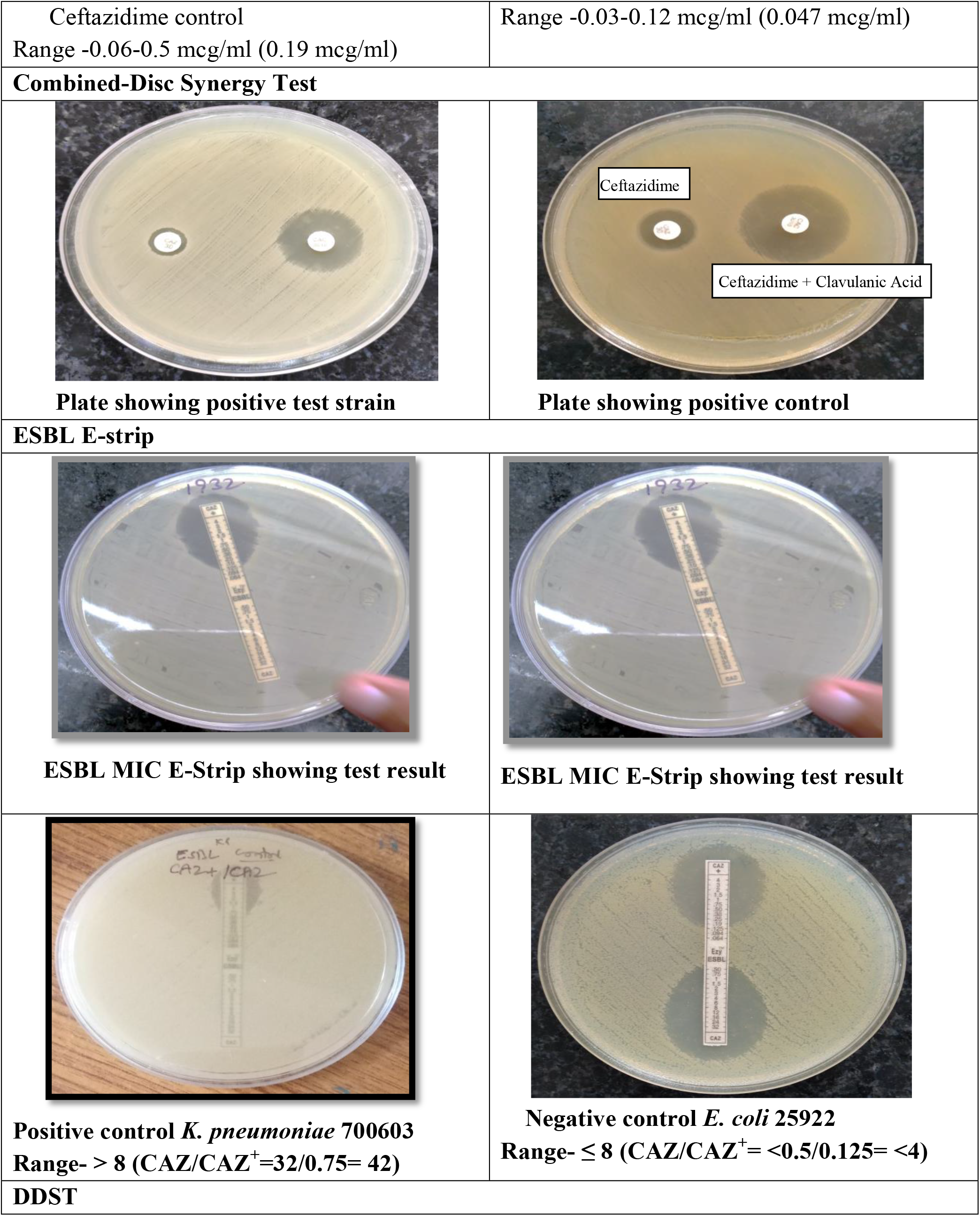
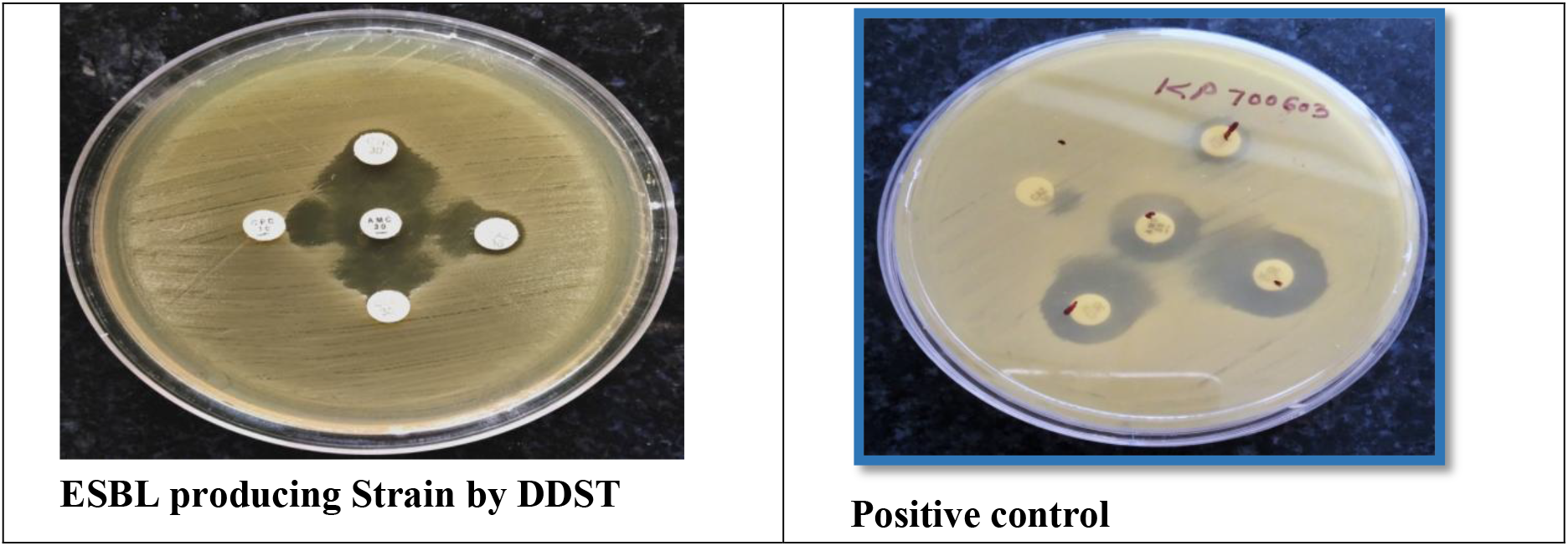
Showing MIC and various Phenotypic test with controls.

Molecular results: The prevalence of bla-CTX-M (80.36%, 45/56) was highest, followed by bla-TEM (50%, 28/56) and bla-SHV (10.71%, 6/56). Among all ESBL positive MDR *E. coli* isolates, two strains (3.57%) were having all the three genes i.e. bla-TEM, bla-SHV and bla-CTX-M. While genes i.e. bla-TEM and bla-SHV were simultaneously present only in three strain (5.36%). bla-TEM and bla-CTX-M were present in twenty five strains (44.64%) and bla-SHV and bla-CTX-M were present in only three strains (5.36%). However, frequency of prevalence of single gene for ESBL was two (3.57%) for bla-TEM and two (3.57%) for bla-SHV, while twenty one (89.29%) for bla-CTX-M. The overall ESBL production among all MDR *E. coli* is 89.29% (50/56) which includes any one of the three genes, Figure-3.

**Figure 3.**
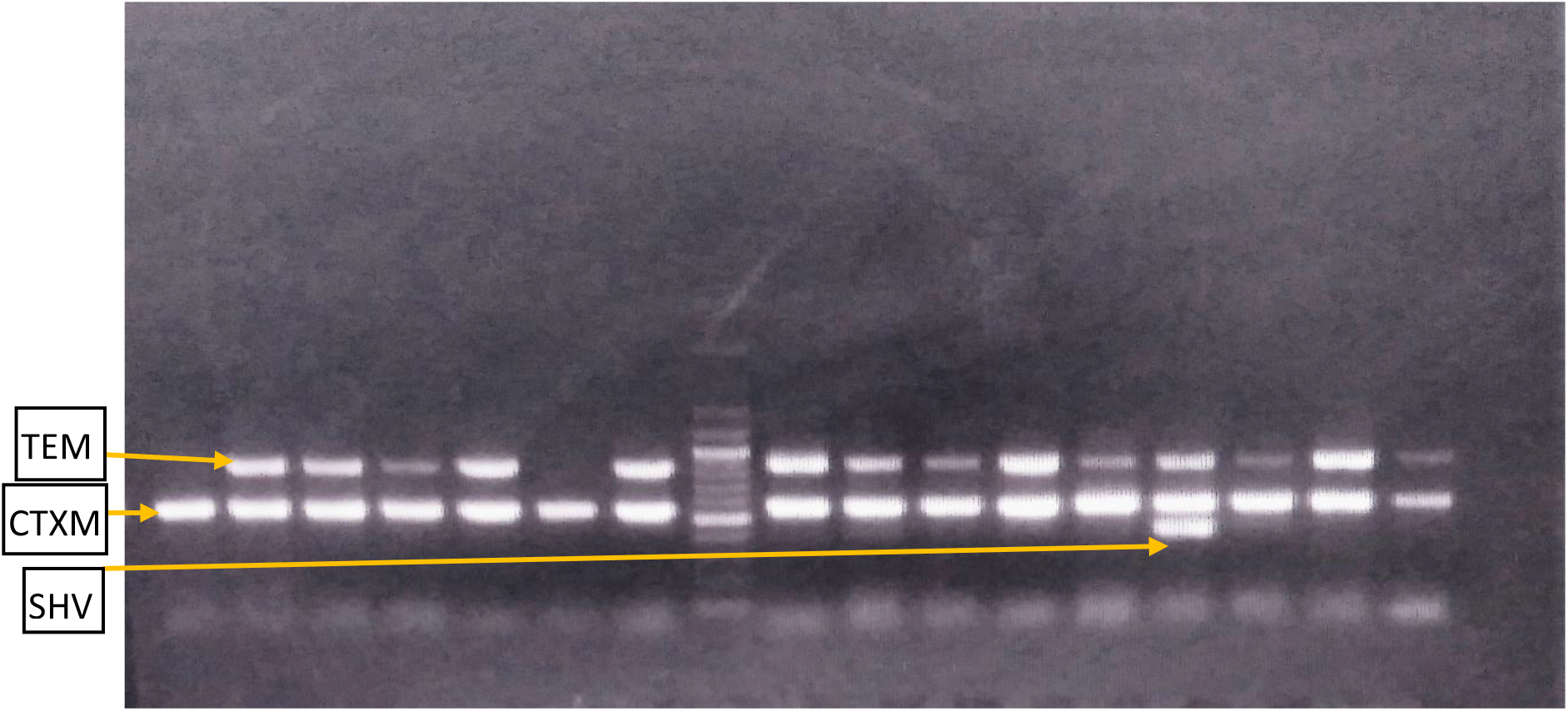
Gel picture showing bands at 867bp for bla-TEM, 550 for bla-CTX-M, and 390 for bla-SHV of PCR products of *E. coli* strains.

## Discussion

Currently Cephalosporins are the drugs of choice for the treatment of infections caused by the Enterobacteriaceae but indiscriminate administration leads to increased resistance to this class of antibiotics (1-Paterson 2005). The emergence of ESBLs producing Gram-negative microorganisms pose significant diagnostic and therapeutic challenge in the management of the infection and impact on the outcome of patient (18-Wong-Beringer 2001) and compromises the activity of wide-spectrum antibiotics including third generation Cephalosporins and other β-lactam antibiotics, Aminoglycosides, Fluoroquinolones and Sulfonamides (19-Hemalatha 2007, 20-Sood 2012, 21-Kothari 2008, 22-Sabharwal 2012 & 23-Dash 2013), thus contributing to become MDR and they can also pass the gene to other clinical strains. So the early detection of these strains and antimicrobial therapy is most important in the treatment. CLSI recommends testing for ESBLs among Gram-negative organisms resistant to 3^rd^ generation Cephalosporins (14-CLSI). Some study reported urine was the major source of ESBL producers (24-Shanthi 2010, 25-Iraj 2010, 26-Saba 2012) while other researcher reported ESBL production was highest in *E. coli* among various GNB isolates (27-Abhilash 2010, 28-Wattal 2010 & 29-Umadevi 2011).

A total 56 patients of UTI were included in this study, of them 76.79% were female, and 23.21% were male, which is in agreement with earlier studies (30-Oladeinde 2011, 20-Sood 2012 &, 31-31-Kashef 2010 & 23-Dash 2013). Short perineum, short urethra and sexual activity has been reported as the factors which influence the higher prevalence in women (32-Omoregie 2008).

The prevalence of MDR *E. coli* isolates isolated from UTI cases was highest within 21 to 30 years of age group (25%), followed by 31 to 40 years (21.43 %).

The prevalence of UTI was highest within 21-30 and 31-40 years of age group i.e. 25.58%, among the female patients which shows that young female patients had highest prevalence rate. This result is in agreement with previous studies (30-Oladeinde 2011, 20-Sood 2012, 33-Shaifali 2012 & 23-Dash 2013). Whereas majority of the isolates among male UTI patients were obtained from age group 21-30 years i.e. 30.77% then followed by 71-80 year age group 23.08%. Elderly males (>70 years) had a higher incidence of UTI (23.08%) when compared with the elderly females (6.98%). Our finding is similar to study conducted by Sood et. al. (20-Sood 2012). This is probably because with advancing age, the incidence of UTI increases among males due to prostate enlargement and neurogenic bladder (34-Das 2006).

Based on antimicrobial susceptibility profile Fosfomycin, Tigecyclin, Cloramphenicol & Nitrofurantoin were the most potent drugs against ESBL producing *E. coli* strains with susceptibility rates of 100%, 80.36%, 58.93% and 53.57% respectively, hence can be used for the treatment of the infection caused by MDR *E. coli*. Meropenem, Tobramycin, Gentamycin, Netilimycin and Imipenem showed moderate activities against these strains with 23.21%, 19.64% and 17.86%, 17.86% and 16.07% susceptibility rates. The highest rates of resistance were observed for Cefotaxime, Ceftriaxone, Ampicillin, Norfloxacin, Levofloxacin and Nalidixic acid i.e. 100% thus treatment with these drugs should not be included in empirical therapy. A study from Hyderabad by 35-Subhalaxmi 2010 showed that only 8% *E. coli* isolates were sensitive to CTR, which is a frequently used empirical antibiotic and in concordance to our study. Due to high resistance to CTX as well as CTR and CAZ, it is suggested that these antibiotics may not be used as a drug of choice for the treatment of infection caused by MDR *E. coli* isolates. The MIC of the isolates that were resistant to third generation Cephalosporins were in the range of 12-256 µg/ml against the Ceftazidime and 32-256 µg/ml for Cefotaxime. 98.21% of the isolates had an MIC 256 µg/ml for Cefotaxime and 96.43% for Ceftazidime. Among 56 MDR *E. coli* isolates 46 drug resistance patterns have been observed in this study and higher resistance was observed in ward patients as compare to out-patients.

In one of the study, the resistances were observed to Co-trimoxazole (82.5%), Ciprofloxacin (80%), Ampicillin (97.5%) and Nalidxic acid (95%) respectively (36-Mukherjee 2013). The other reports from different geographical locations in India revealed a high resistance against a majority of the antibiotics such as Co-trimoxazole, Gentamycin, Ciprofloxacin, and Amikacin (37-Peripi 2012). Ghotaslou et. al. reported high rates of resistance were observed against Ampicillin, Co-trimoxazole and Ciprofloxacin prevalence of ESBLs rate among *E. coli* was 38.3% (38-Ghotaslou 2018). Mukherjee et. al. reported that the ESBL production was 45% in the *E. coli* isolates (36-Mukherjee 2013). The prevalence of ESBL producing Gram-negative isolates was reported in various hospitals in India in the range of 19-60% (39-Bhattacharya 2011).

As we have selected 3^rd^ generation Cephalosporin resistant isolates which means that almost all the isolates would need further testing for ESBL detection. The Health Protection Agency recommends testing of Cefpodoxime or both Cefotaxime and Ceftazidime as a first screening test (40-Health Protection Agency 2008). The recommended approach for ESBL detection is to use a screening test then a confirmatory method for ESBL detection on all 3^rd^ generation Cephalosporin strains (14-CSLI 2020).

The ESBL production by phenotypic methods was observed in 100%, 94.64% and 67.86% by E-strip method, DDST and CDST in MDR *E. coli* isolates. We have found that prevalence of ESBL production among hospitalized patients and out patients by E-strip, CDST and DDST were 100%, 72.72% (24/33) & 100% and 100%, 60.86% (14/23) & 86.95% (20/23) which is similar with findings of other investigator (41-Vemula 2011, 27-Abhilash 2010, 42-Sharma 2013). The difference observed in detection of ESBL positive by two different methods may be justified by the lower sensitivity of different phenotypic methods and the influence of environmental factors on the incidence of resistance (43-Yazdi et. al. 2012, 44-Ravi et. al. 2011). In CDST the sensitivity is low probably due to Ceftazidime alone used for the detection of ESBL.

Sageerabanoo et. al. from Puducherry has reported 77.02% *E. coli* isolates as ESBL producer, previously screened as third generation Cephalosporin resistant (45-Sageerabanoo 2015). Sharma et. al. 2013 reported 57.18% strains of *E. coli* were ESBLs producer by disc diffusion test (42-Sharma et al. 2013), while Grover et al from Delhi reported 56.67% strains of *E. coli* ESBL producer (46-Grover 2017).

One of the study from Jaipur reported 57% ESBL production in *E. coli* (42-Sharma 2013). The incidence of ESBL production in various studies reported in India varies from 60% to 80% (18-Wong-Beringer 2001), followed by China (60%), East and South-east Asia (<30%) and other countries such as Europe, Australia and North America (range 5-10%) in among *E. coli*. A steady rise has been reported in the prevalence of ESBL producing *E. coli* in India, i.e. 18-40% during 2003 to 2008 while it rose to 40-75% during 2009 to 2012 (47-Livermore 2012, 48-Tankhiwale 2004, 49-Taneja 2008, 50-Hawser 2009). Different studies shows ESBL production rate varies from 17-70% in India (51-Gopalakrishnan et. al. 2010, 24-Shanthi & Sekar 2010, 41-Vemula & Vadde 2011, 52-Mubarak et. al. 2011, 53-Ali et. al. 2004, 54-Patel et. al. 2012).

In the present study, one isolate was ESBL non-producer by E-strip and DDST methods while showing ESBL production by CDDT and PCR (showing SHV gene). Another strain of MDR *E. coli* which is ESBL non-producer by DDST and CDDT method but surprisingly ESBL producer by E-strip and PCR (bla-CTX-M gene). We have also observed that results of DDST and E-strip method have almost similar sensitivities although the former being the relatively cheaper method can be used effectively in hospital setting.

The data reported from neighboring counties for ESBL production among *E. coli* was 50% from Turkey and 72% from Pakistan respectively (55-Senbayrak Akcay 2014, 56-Qureshi 2013, 42-Sharma 2013). Nagdeo et. al. reported 78.22% *E. coli* as ESBL producer (57-Nagdeo 2012).

PCR has been considered to be a reliable method for detection of ESBL compared to DDST and E-strip method (58-Grover 2006). Furthermore, phenotypic tests for ESBL detection only confirm whether an ESBL is produced but cannot detect the ESBL subtype and furthermore definitive identification is only possible by molecular detection methods (43-Yazdi 2012).

### Molecular

There are so many types of ESBLs genes like TEM, SHV, CTX-M, OXA and AmpC etc. found in *E. coli*. The bla-CTX-M was the commonest gene present in strains i.e. 80.36% followed by bla-TEM 50%, and bla-SHV 10.71%. Similar results have been obtained in many studies in India and other countries for bla-CTX-M gene prevalence (59-Krishnamurthy 2013, 60-Al-Algamy et. al. 2009) but in case of bla-TEM and bla-SHV, it is contradictory. In a study conducted from 10 different Indian sites by Welsh et al. (61-Welsh 2006), bla-CTX-M was the most common genotype isolated. Similar results have been reported in Europe, Latin America and other countries (62-Dhillon 2012).

Similarly Mubarak et. al. reported emergence and dissemination of 87% CTX-M-15 Producing *E. coli* in the UAE but the SHV gene was detected in 13% of the strains by them which is similar to the present study (52-Mubarak 2011).

Bali et. al. reported TEM in 72.72%, SHV in 9.09% and 22.72% CTX-M encoding genes were reported in *E. coli*, which is discordant from present study (63-Bali 2010), while Sharma and Vaida reported CTX-M genes i.e. 80% and 96% in *E. coli* which is concordant to our study (42-Sharma 2013 and 64-Vaida 2010). Some of the studies states that the CTX-M gene is the most prevalent ESBL encoding gene worldwide and is replacing TEM and SHV type in European and Asian countries, which is in concordance to the present study (65-Livermore 2007, 66-Bonnet 2004, 67-Naseer 2011).

A study from Thailand reported 99.6% of ESBL producing *E. coli* isolates carried out bla-CTX-M (68-Pattarachai 2008). The bla-TEM and bla-SHV groups were detected in 77.0% and 3.8% ESBL producing *E. coli*. Another study from Turkey by Nazik et. al. reported the most frequent β-lactamase type was CTX-M (92%), followed by TEM (39%), SHV (5%) (69-Nazik 2011).

The ESBL producers usually carry multiple resistant plasmid genes which may be related to complex antimicrobial resistance (70-Ye QH 2011) and it has also been reported in our study, two isolates of MDR *E. coli* had all three genes (3.57%%) bla-CTX-M, bla-TEM and bla-SHV, twenty five (44.64%) isolates had two genes i.e. bla-CTX-M and bla-TEM, three isolates (5.36%) had two genes i.e. bla-CTX-M & bla-SHV and three isolates (5.36%) of MDR *E. coli* had bla-TEM and bla-SHV. Similar results have been reported by other researchers (59-Krishnamurthy 2013, 43-Yazdi et. al. 2012, 71-Yuan et al 2012). According to Goyal et. al., 57.3% of strains harbored 2 or more ESBL genes (72-Goyal 2009), while Bali et. al. observed that 19.2% ESBL isolates carried more than one type of β-lactamase genes (63-Bali 2010). 54.29% isolates in our study carried more than one type of β-lactamase genes. Sharma et. al. observed 77.5% isolates carried more than one type of genes, which is quite high as compared to our study and previous studies, Sharma et. al. also reported that 32.5% isolates harboring all the three β-lactamase genes, 22.5% isolates harboring bla-TEM and bla-CTX-M genes, 15% isolates harboring bla-SHV and bla-CTX-M genes and 7.5% isolates harboring bla-TEM and bla-SHV genes (42-Sharma 2013).

## Conclusion

It is unfortunate that all MDR *E. coli* isolates included in this study was found to be resistant to most of the routine antibiotics including third generation Cephalosporins, thus reducing the clinical utility of these drugs. The resistance patterns of antibiotics, frequently used for microorganisms vary over the time, so the empiric antibacterial therapy should be based on a local susceptibility and resistance profile. Incorrect assessment of antibiotic resistance may lead to inappropriate antibiotic prescription, which in turn may direct bacteria to produce new resistance genes by selective pressure. In our study, expressions of various β-lactamases either singly or in combination are in agreement with the other studies. The prevalence of ESBL production limits the clinical use of β-lactams. Our findings suggest that the presence of bla-CTX-M as the predominant genotype in *E. coli* isolates in Delhi. In view of emerging AMR the practice of routine ESBL testing along with antibiogram would be useful for all cases which will help in the proper treatment of the patient and also prevent further development of bacterial drug resistance.

